# Landscapes and dynamic diversifications of B-cell receptor repertoires in COVID-19 patients

**DOI:** 10.1101/2020.12.28.424622

**Authors:** Haitao Xiang, Yingze Zhao, Xinyang Li, Peipei Liu, Longlong Wang, Meiniang Wang, Lei Tian, Haixi Sun, Wei Zhang, Ziqian Xu, Beiwei Ye, Xiaoju Yuan, Pengyan Wang, Ning Zhang, Yuhuan Gong, Chengrong Bian, Zhaohai Wang, Linxiang Yu, Jin Yan, Fanping Meng, Changqing Bai, Xiaoshan Wang, Xiaopan Liu, Kai Gao, Liang Wu, Longqi Liu, Ying Gu, Yuhai Bi, Yi Shi, Shaogeng Zhang, Chen Zhu, Xun Xu, Guizhen Wu, George F. Gao, Naibo Yang, William J. Liu, Penghui Yang

## Abstract

Severe acute respiratory syndrome coronavirus 2 (SARS-CoV-2) has caused the pandemic of coronavirus disease 2019 (COVID-19). Great international efforts have been put into the development of prophylactic vaccines and neutralizing antibodies. However, the knowledge about the B cell immune response induced by the SARS-CoV-2 virus is still limited. Here, we report a comprehensive characterization of the dynamics of immunoglobin heavy chain (IGH) repertoire in COVID-19 patients. By using next-generation sequencing technology, we examined the temporal changes in the landscape of the patient’s immunological status, and found dramatic changes in the IGH within the patients’ immune system after the onset of COVID-19 symptoms. Although different patients have distinct immune responses to SARS-CoV-2 infection, by employing clonotype overlap, lineage expansion and clonotype network analyses, we observed a higher clonotype overlap and substantial lineage expansion of B cell clones during 2-3 weeks of illness, which is of great importance to B-cell immune responses. Meanwhile, for preferences of V gene usage during SARS-CoV-2 infection, IGHV3-74 and IGHV4-34 and IGHV4-39 in COVID-19 patients were more abundant than that of healthy controls. Overall, we present an immunological resource for SARS-CoV-2 that could promote both therapeutic development as well as mechanistic research.

## INTRODUCTION

The current outbreak of SARS-CoV-2 infection is threatening global public health ^[1]^. The scale of the humanitarian and economic impact of the COVID-19 is driving intense efforts to develop vaccines and neutralizing antibodies (NAb) against COVID-19. Therefore, understanding the principles of the B-cell responses during SARS-CoV-2 infection is of substantial importance for anti-viral vaccine and NAb developmeent.

The B-cell receptors (BCRs) are immunoglobin molecules located on B-cell surfaces to recognize and bind foreign antigens. Upon encountering their specific antigen, B-cells become activated, proliferate, and may differentiate to produce short-lived effective antibody-secreting plasma cells or long-lived plasma cells and memory cells. At the same time, BCRs undergo a process of affinity maturation, which repeats cycles of somatic hypermutation of BCRs and subsequent clonal selection leads to increased binding affinity. BCRs are composed of two heavy chain molecules and two light chain molecules, encoded in humans by the immunoglobin heavy chain (IGH) gene and light chains (IGL or IGK) gene, respectively. Comparison of BCR sequences among individuals is of great interest because repertoires may have similar features if individuals are exposed to the same pathogen, giving rise to convergent antibodies.

Antibody specificity is largely determined by the IGH gene sequence used by each B-cell ^[2]^. The recent developments in high-throughput sequencing have made it feasible to character IGH repertoire in large numbers of samples and it is increasingly being applied to gain insights into the humoral responses in healthy individuals and a wide range of diseases. This technic has also led to advances in our understanding of how the antibody repertoire changes in response to perturbation arising from an initial viral infection, viral evolution, and vaccination. BCR repertoire analysis has been applied, for example, to influenza virus ^[3][4]^, human immunodeficiency virus ^[5]^, varicella-zoster virus ^[6]^ and dengue virus ^[7]^. However, the dynamics of antibody response elicited by SARS-CoV-2 infection remain to be determined.

To understand how B-cell immune repertoire changes over time during SARS-CoV-2 infection, we obtained IGH repertoires from the peripheral blood samples which were collected multiple times from five COVID-19 patients. We classified sequences into clones by lineage clustering analysis, and tracked changes in the unique number of CDR3, Shannon index, the number of high-frequency clones, cumulative frequency of the top 100 clones, and V, J-gene segment usage at different course of time. We also conducted clonotype overlap, lineage expansion, and CDR3 sequence network structure to explore the similarity and varying trends of IGH repertoire status at different time points. Overall, during the high degree of heterogeneity in B-cell clonal dynamics among patients, key common patterns were observed, i.e. the higher clonotype overlap and substantial lineage expansion of COVID-19 patients during 2-3 weeks of illness. The findings in this study provides basic knowledge support for the immune response research of COVID-19 patients and the development of corresponding antibody drugs and vaccines.

## MATERIALS AND METHODS

### Study approval and samples

This study has been approved by the Research Ethics Committee of The Fifth Medical Center of PLA General Hospital, Beijing, China. (approval number: 2020034D), written informed consent was regularly obtained from all patients. Five patients with newly diagnosed COVID-19 and three healthy donors were enrolled. According to the Diagnosis and Treatment Protocol for Novel Coronavirus Pneumonia (Trial Version 4, Released by National Health Commission & State Administration of Traditional Chinese Medicine on January 27, 2020), as blood samples were tested positive for SARS-Cov-2 virus, all patients were diagnosed as COVID-19 patients, with the 85-year-old patient developed severe symptoms, and the other four patients aged between 15 to 45 developed mild symptoms. Blood samples were collected at multiple points during hospitalization. Detailed clinical information is shown in Table 1. The laboratory characteristics were listed in Supplementary Table 1.

**Table 1.**
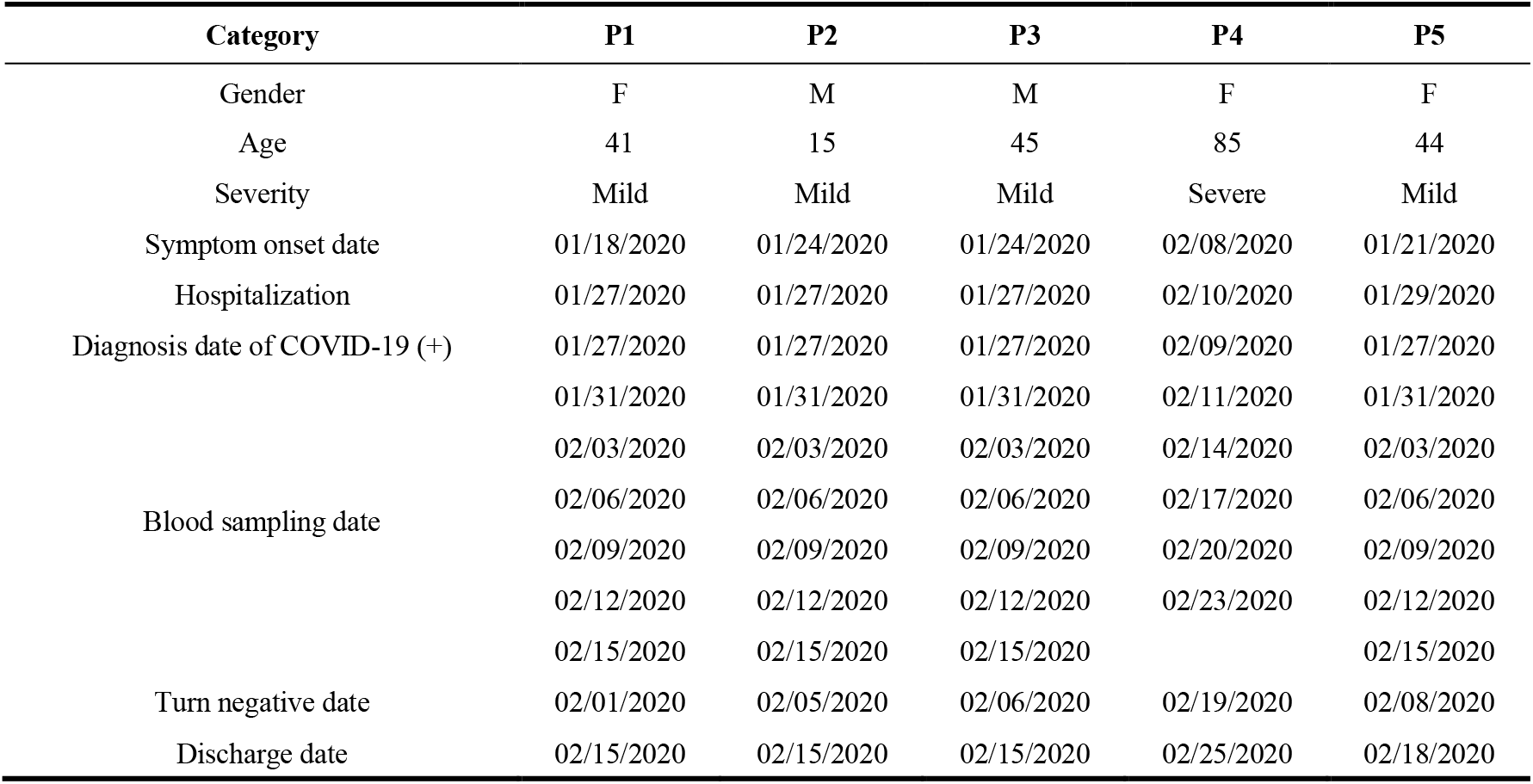
Summary of the clinical information of the 5 convalescent patients.

### Library preparation and repertoire sequencing

Human peripheral blood mononuclear cells (PBMCs) were isolated from blood samples of five COVID-19 patients as well as three healthy donors by density gradient centrifugation in Ficoll-Paque Plus buffer (GE Healthcare, USA). Total RNA was extracted from approximately 1×10^5^ PBMCs of each sample using Trizol Reagent (Invitrogen, USA). cDNA was synthesized with SuperScript II reverse transcriptase (Invitrogen, USA) and Oligo-dN6 random primer. The variable domain of IGH was amplified by using six V segment specific primers designed from FR1 region and one J segment specific primer designed from FR4 region ^[8]^. The PCR program began with an initial denaturation at 95 °C for 15 min, followed by 28 cycles of denaturation at 94 °C for 30 s, annealing of primer to DNA at 60 °C for 90 s, and extension by 2 × multiplex PCR master mix (QIAGEN, USA) at 72 °C for 30 s. Following the cycling, the reaction was at 72 °C for 5 min, with a final step at 12 °C to hold the terminated reaction. The amplified PCR products were then purified by the same volume of the Ampure-XP beads. Subsequently, the purified PCR products were end-repaired and ligated with a sequencing adapter. The sequencing libraries were quantified using Qubit ssDNA HS (High Sensitivity) Assay Kit (Invitrogen, USA) and sequenced by MGISEQ-2000 (MGI, China) platform with 400 bp single-end reads.

### Basic data processing

The single-end 400 bp reads of the samples from the five COVID-19 patients sequenced at MGISEQ-2000 platform were analyzed by IMonitor ^[9]^ software. Briefly, the raw reads were filtered and trimmed by SOAPnuke ^[10]^ to remove low-quality reads and adapter sequences. Reference germline gene segments were collected from IMGT database ^[11]^. BLAST ^[12]^ program was used to align all clean reads to a reference, and then a second alignment procedure was executed to improve the alignment accuracy. Subsequently, structure analysis was performed to deduce the structure of BCR molecular, including V, D, and J segments usage, the nucleotide and amino acid sequence of the CDR3 region, and the deletions and insertions at rearrangement sites. Finally, several characters reflecting the status of the immune repertoire were counted, such as V-J pairing, V/J usage, CDR3 sequence frequency, and CDR3 length distribution.

### Lineage identification and effective data normalization

We define sequences belonging to the same clonal lineage as those sharing the same variable (V) and joining (J) germline gene combination, CDR3 length, and at least 90% nucleotide sequence identity in the CDR3 region. According to this definition, we first grouped sequence with identical V and J germline gene origination and same CDR3 length, following clustering those group by CD-HIT (version 4.6) ^[13]^ with the following parameters: -c 0.9, -G 1, -b 20, -d 0 and -n 9. To eliminate the impact of the difference in data volume on clonal diversity, we normalized each of the IGH clones to a unanimous number. Firstly, we counted an effective data size (with an “in-frame” tag in the IMonitor result) for each sample and removed some samples with significantly fewer effective data than others. Secondly, normalized data with the same size were extracted randomly from each sample by an in-house Perl program.

### Feature parameters analysis and comparison of usage frequency of V/J germline gene segments

There are several commonly used parameters reflecting the features of the BCR repertoire, including unique CDR3 number, Shannon index, high-frequency clone number, and cumulate frequency of the top 100 clones. we are using an in-house Perl script to count those parameters base on normalized data from each sample. The Shannon index is frequently used as measures of repertoire diversity, we calculated the Shannon index of observed clones from normalized data using the following function:

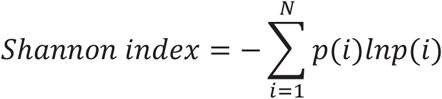

where N is the total number of unique CDR3, and p(i) is the frequency of a single CDR3 in the BCR repertoire.

After data normalization, gene usage frequency of V and J segments were counted and a k-means hierarchical clustering process was executed by R package “hclust” to compare V/J segment usage similarity both within and among individuals.

### Overlap analysis of BCR repertoire

Repertoire overlap analysis is a similarity measure to evaluate the correlation of overlapping clonotypes between samples. The Jaccard index, also known as intersection over union and the Jaccard similarity coefficient, can be used to gauging the similarity and diversity of different repertoires. To explore the impact of onset time on the status of the BCR repertoire, we counted the Jaccard index to assess the association between the repertoire at different onset times, the index is calculated as follows:

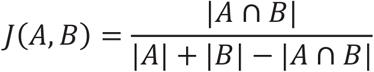

Where |A | denotes the total number of unique CDR3 in sample A, |A∩B| represents the intersection of the unique CDR3 number in samples A and B.

We calculate pairwise overlap rate as Jaccard index multiplied by 100, and divided the pairwise overlap rate into 3 clusters: the first cluster is “2-3 week adjacents” including data of time points 12-13 days versus 15-16 days and 15-16 days versus 18-19 days after onset; the second cluster is “other adjacents” which contains two adjacent time points except “2-3 week adjacents”; the third cluster is “non adjacent” which contains all of the paired non-adjacent time points. Two-sample Wilcoxon test (Wilcoxon test in R) was used to compare the difference between clusters.

### Lineage expansion analysis

A lineage expansion analysis was performed based on the normalized data, Firstly, a join samples process was executed to join lineage information at all time points inside each patient together, normalized lineage frequency was computed as the geometric mean of lineage frequencies that comprise a given joint lineage in intersected time points if the lineage is missing, its frequency is set to 1e-9, only the most abundant CDR3 sequence was selected as a representative clonotype in each lineage. Secondly, an R program originally generated by VDJtools ^[14]^ with some modifications was used to display the expansion and contraction of lineages.

### CDR3 sequence network generating

The network is generating through the BCR CDR3 amino acid similarity, where each vertex in the network represents a CDR3 amino acid sequence, the size of the point represents the abundance of the clone, any two vertices meet the following three conditions, including same length, same V and J gene usage, and only one amino substitution are connected with an edge. The diagram was plotted by the “igraph” R package (https://igraph.org/redirect.html).

### Parameters of the network calculation

The parameters of the network, including vertices, cluster number, and diameter of the largest cluster, are calculated using “V”, “clusters”, “diameter” functions in the “igraph” package. The Gini coefficient is used as measures of inequality in distribution, we calculated the Gini coefficient of clone frequency distribution and cluster clone number distribution using the following function:

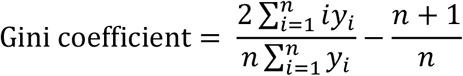

where y(i) is the frequency of a single clone, and n is the total unique clone number in the calculation of clone frequency Gini coefficient; y(i) is the clone number of a single cluster, and n is the total cluster number in the calculation of cluster Gini coefficient.

## RESULT

### Basic Clinical Characteristics of COVID-19 patients

Among the five patients diagnosed with COVID-19, the 85-year-old patient developed severe symptoms, and the other four patients aged between 15 to 45 developed mild symptoms (Table 1). Blood samples were collected at day 2-5 after diagnosis with 2-3 days intervals until they were tested negative for SARS-CoV-2 on day 17-19. All of them were discharged from the hospital upon recovery. Notably, 3 of the 5 patients are from the same family, that is, patient 2 is son of patient 1 and patient 3.

During the process of COVID-19 diagnosis, several clinical indicators were detected by routine blood examination. At least three of the 5 patients (2 of them have too few tested indicators and can’t be reasonably counted) showed pronounced lymphopenia. Along with the progression of COVID-19, the percentage of T lymphocytes went through a process of decreasing firstly and then increasing. The phenomenon of lymphopenia in COVID-19 patients in our results is highly consistent with previous studies. It is well established that the lymphopenia is a major immunological abnormality occurs in COVID-19 patients ^[15, 16, 17, 18]^. However, we found that the percentage of B lymphocytes showed an opposite trend to T cells, among these patients, the percentage of B cells in patient P3, P4 and P5 reached the highest value until 14^th^, 6^th^ and 17^th^ day after the onset of the disease, respectively (**Supplementary Table 1**). At the same time, B cell clonal expansion reached the peak value from immune repertoire analysis (**Fig 3**).

### IGH repertoire altered substantially upon infection with SARS-CoV-2

We performed longitudinal IGH repertoire sequencing to examine the global effects of SARS-CoV-2 infection on the B-cell compartment (**Fig 1 a)**. To enable direct comparisons among time points of each patient, datasets of all time points of each patient were sub-sampled to a unanimous number of clonotypes. We investigated the diversity of the IGH repertoires through four parameters, which are unique CDR3 number, Shannon index, high-frequency clone number (frequency > 0.1%), and cumulate frequency of top 100 clones with the highest frequency. These parameters fluctuate substantially throughout time with no common tendencies among patients (**Fig 1 b-e**), which may due to the dramatic change of the predominant clones and lineages.

**Figure 1.**
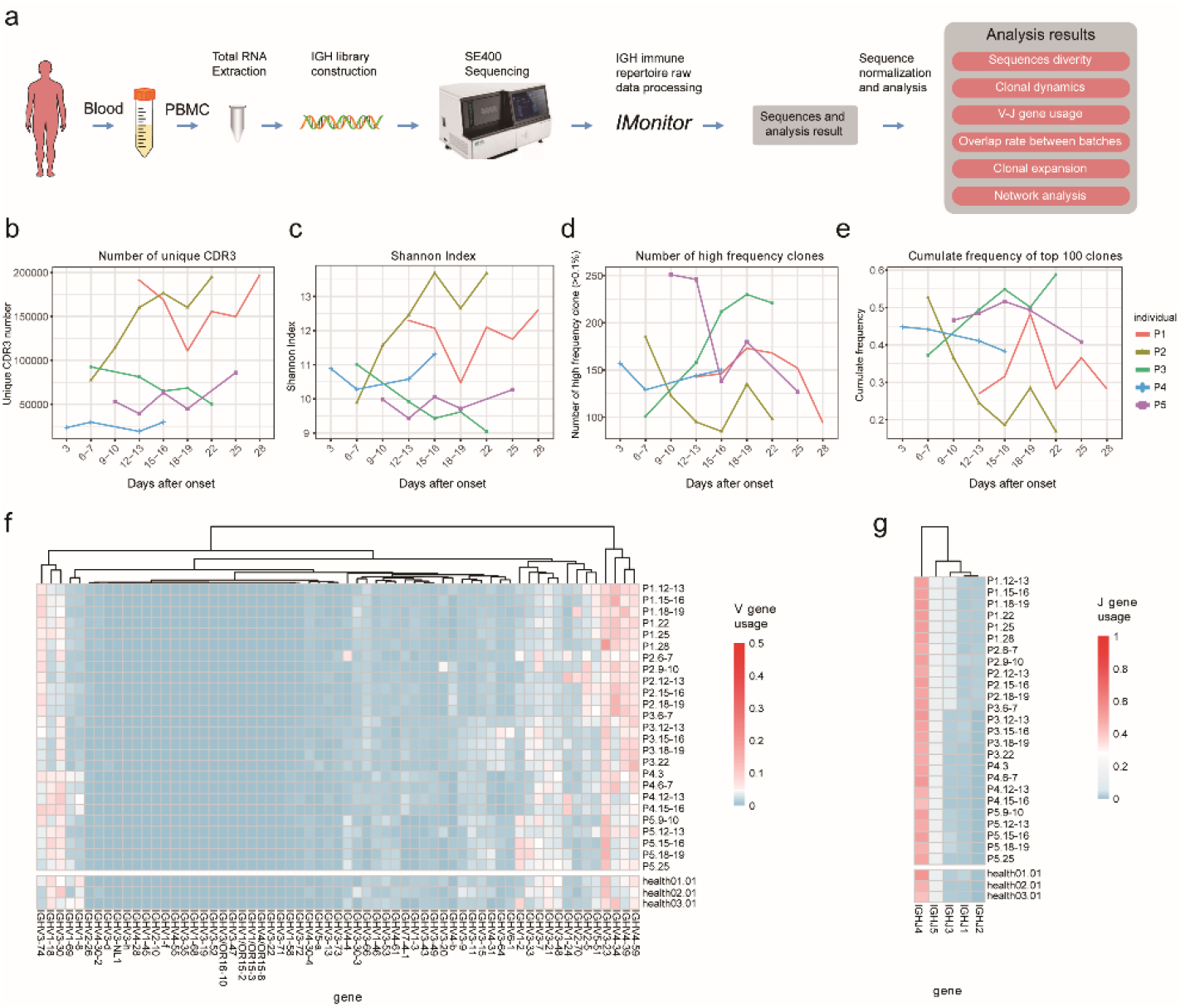
Schematic diagram of the experimental process and overall dynamic changes of the BCR repertoires. **a.** Schematics of the experimental design and bioinformatic workflow. **(b–e)** Unique CDR3 numbers, Shannon index, high frequency (>0.1%) clone numbers and the cumulative frequency of the top 100 clones of the BCR repertoire. Each dotted line represents the dynamic tendency of in a single sample over time after onset. **(f-g)** Heat map shows the gene usage proportion inside each sample for the V/J genes listed at the bottom on the figure, each row corresponds to an individual, each column corresponds to a V/J germline gene, the color in each tile corresponds to frequency of V/J gene in the repertoire, the deeper the color, the higher the frequency.

Next, we thought to test whether there are preferences of V and J gene segments during SARS-CoV-2 infection. By using hierarchical clustering strategy, we compared the V and J gene usage frequency of COVID-19 patients with that of healthy donors. However, compared with healthy donors, we did not find overall preference in the usage of V or J genes among COVID-19 patients, the overall use of each V and J component was highly nonuniform within each individual (**Fig1 f, g**). The most frequently observed V segments were IGHV3-23, IGHV3-30, IGHV4-34, IGHV4-39, and IGHV4-59, which is similar to previous studies ^[19, 20]^. The IGHV3-74, IGHV4-34 and IGHV4-39 in COVID-19 patients showed more abundant than that of healthy controls. Preferred J gene segments showed a high degree of consistency among all time points in both patients and healthy donors (**Fig 1 g**), with the most frequent J segment is IGHJ4, this is consistent with previously report ^[21]^. For each individual, the frequency of distinct gene segments usage over the infection time course was fairly stable, shown as high correlations in the heatmap of **Fig 1 f** and **g**, suggesting that overall V, J segment use is not markedly altered by SARS-CoV-2. This phenomenon may reflect that the human immune system is highly variable between individuals but stays relatively stable over time within a given person ^[22]^.

### Repertoire overlap analysis revealed higher similarity of immunological status between the adjacent time points in 2-3 weeks since symptom onset

To better understand the potential mechanism behind changes in IGH repertoire diversity, we examined how the composition of the clonotype changed during the infection. We evaluated the level of clonotype overlap between any two time points of each patient by using an all-versus-all pairwise overlap analysis. In general, the overlap rate between different time points ranges from 0.36% to 6.57% (average 2.13%) in each individual (**Fig 2 a-e**). As expected, samples from adjacent sampling batches share more clones than those from other batches. A notable rule that shows in almost each patient is that the overlap rate of adjacent batches peaks at around 2-3 weeks. We found that “2-3 week adjacents” cluster shared significantly more clones than “other adjacents” (p=7.78 x 10^−3^) and “non-adjacent” (p=4.52 x 10^−5^). (**Fig 2 f**). This indicates that, compared with other time points, the composition of the IGH immune repertoire within individuals is more similar during 2-3 weeks after the onset. It is highly likely that 2-3 weeks after onset is a critical stage of B-cell immune response.

**Figure 2.**
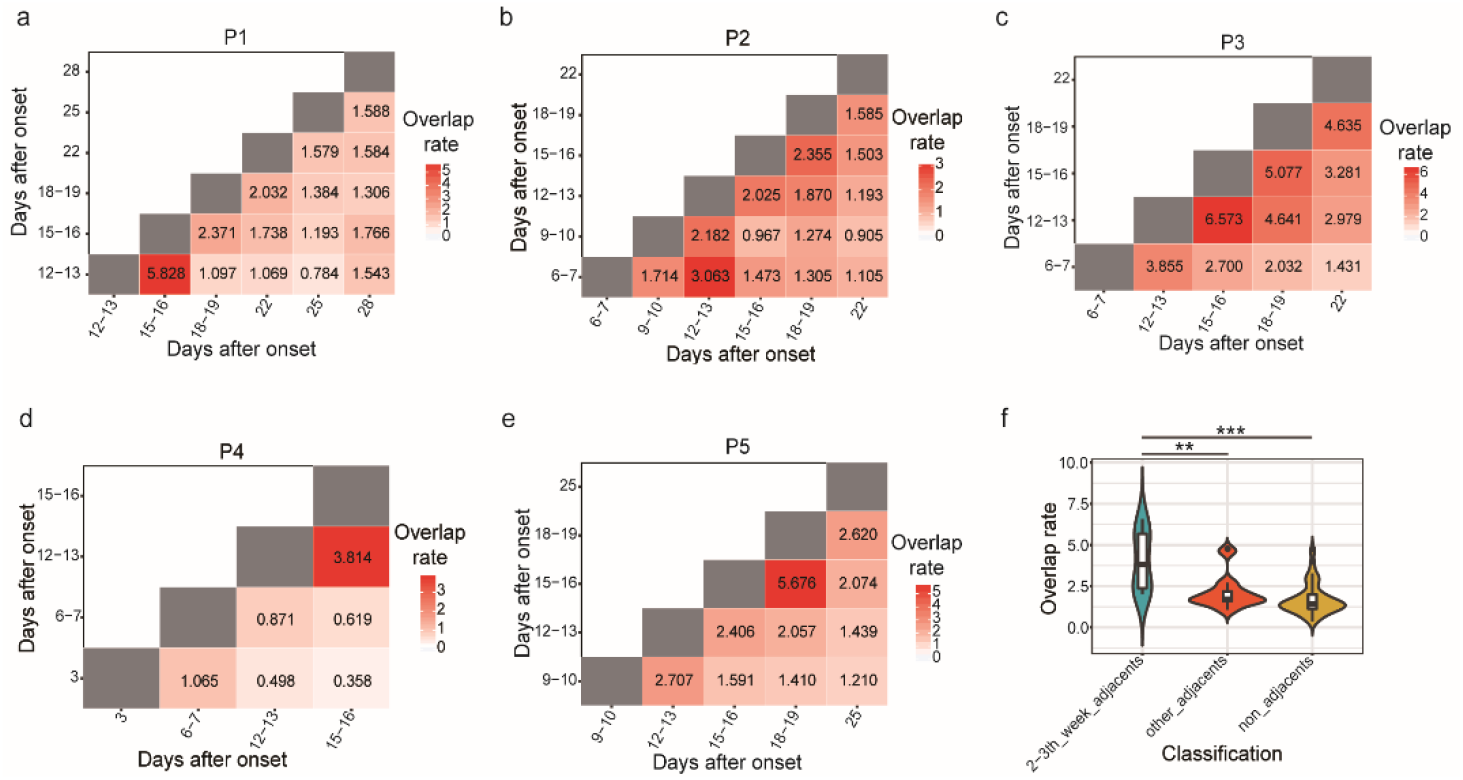
Overview of BCR repertoire overlap changes among various time points after onset. **(a-e)** Each sub-graph is a heatmap describe an all-vs-all intersection between two time points in a single patient, the value in each tile represents the clonotype overlap rate between two time points as shown on the X-axis and Y-axis, the color in each tile corresponds to clonotype overlap rate between two time points, the deeper the color, the higher the overlap rate. f. The comparison of clonotype overlap rate among various time point categories. Comparison were performed using paired Wilcoxon test. ** P < 0.01, *** P < 0.001.

### Tracking analysis revealed evidence of lineage expansion of IGH repertoire during SARS-CoV-2 infection

Next, we investigated the relative degree of clonal expansion. To measure the alterations in lineage frequency over time, we first defined clonotypes as a lineage if they meet 3 criteria: sharing the same VH and JH germline gene segments, having the same nucleotide length in the CDR3 region, and at least 90% nucleotide identity in the CDR3 region. Using this strategy, we computed the geometric mean of linage frequencies of all samples and performed a lineage tracking analysis using VDJtools. Here, we defined the highly expanded lineages as the most 100 abundant lineages (top 100 lineages). We observed an evident lineage expansion process over time. As shown in **Fig 3 a-e**, the cumulative frequencies of the top 100 lineages of almost all patients increased after the second or third weeks following the onset of illness, with the cumulative frequencies of the top 100 lineages peaked at day 18-19 in Patient 1 and Patient 5, day 12-13 in Patient 3, day 6-7 in Patient 4 and both peaked at day 6-7 and day 18-19 in Patient 2. After that, this part of lineages began to decrease gradually. This result indicates that during the 2-3 weeks of disease progression, the patient’s IGH repertoire experienced a substantial lineage expansion.

**Figure 3.**
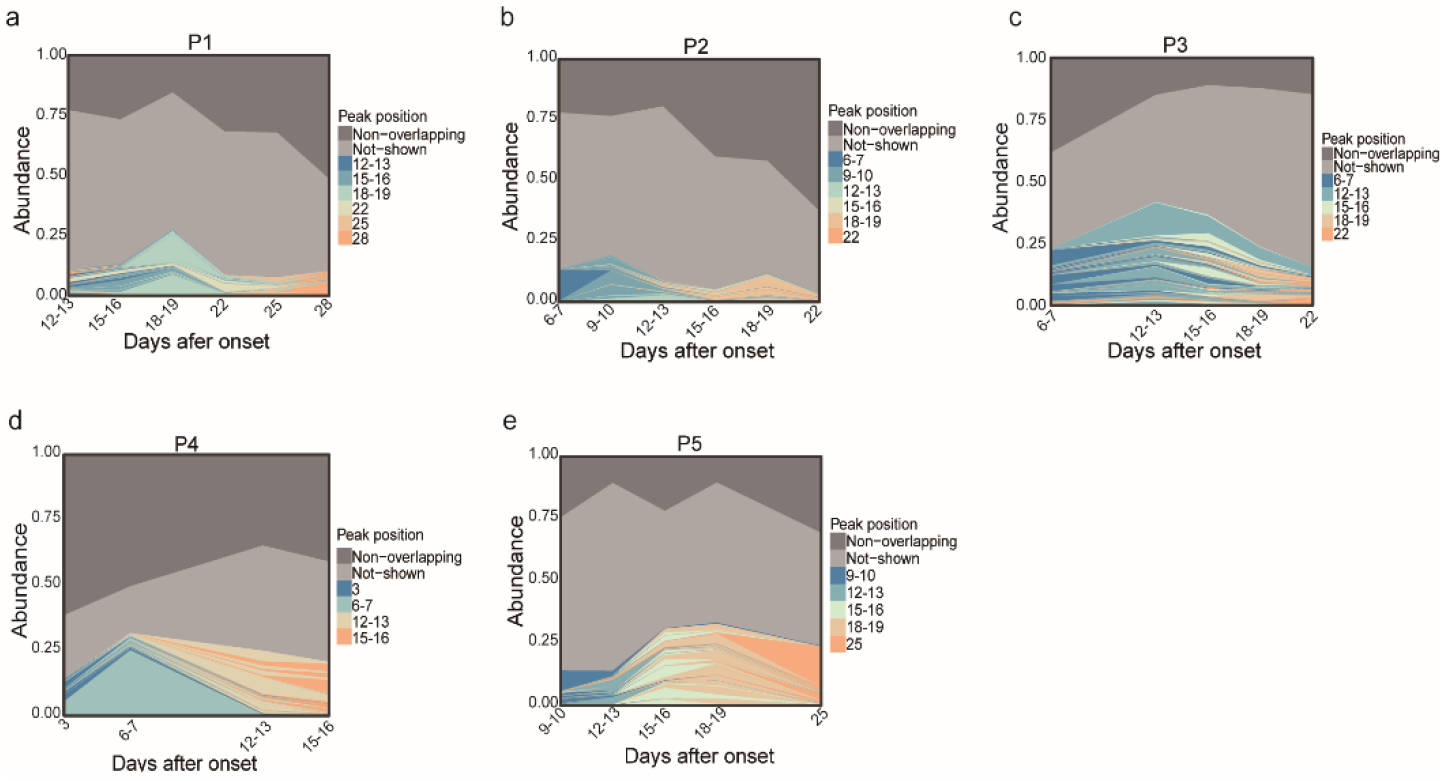
BCR lineage expand and contract over time after COVID-19 onset. Each sub-graph shows the detailed profiles for lineage expansion/constriction in a single patient. Top 100 lineages are colored by the peak position of their abundance profile, other lineages in each BCR repertoire are divided into two groups: non-overlapping (“Non-overlapping”) lineages are marked in dark gray, and the remaining lineages (“Not-shown”) are marked in light gray.

### Network Features delineate the evolution of BCR repertoire landscape

BCRs with similar CDR3 amino acid sequences are supposed to be gathered in clusters based on antigen-driven somatic hypermutation (SHM). To investigate the degree of clone convergence during the disease, we build the BCR repertoire CDR3 sequence network for each individual using data of different times after the onset of the disease based on a specific approach ^[23]^. Networks of all the 5 patients after symptom onset have been plotted in **Supplementary Fig 1** and the network of the 41-year-old female patient 1 (P1) at different time point is demonstrated in **Fig 4a** since its relatively high number of CDR3 clones (vertices). Here, a clear clonal clustering phenomenon could be seen from the network of this patient on day 18-19 (**Fig 4a**). Subsequently, three network-related parameters, namely diameter of the largest cluster, Gini coefficient of cluster clone number, and Gini coefficient of clone frequency, were counted and summarized in **Fig 4 b-d**. There was a similar trend that first goes up and peaks around 2-3 weeks and fall afterward in these three parameters of almost all patients. Each parameter of the network has its implication, firstly, the diameter parameter exhibit in **Fig 4b** is the length of the longest geodesic (edges between any two reachable vertices) of a network. The result indicates higher evolution level of BCR clones in the repertoire from 2-3 weeks. Secondly, the Gini coefficient of cluster clone number which represents the level of clonal clustering also peaked around 2-3 weeks. Thirdly, the Gini coefficient of clone frequency which indicates the skewed expansion of several clones were very much synchronized with the former parameter, despite they were relatively independent. Moreover, the most abundant clones usually don’t appear in the largest cluster (**Fig 4d, Supplementary Fig 1**), the formation of big clusters may be due to the rapid iteration of naïve B-cells to generate affinity maturation, while the clonal expansion may result from the proliferate of the effector cells. Nevertheless, both the cluster clone number increasement and clonal expansion are supposed to be triggered by the reaction to counteract the SARS-CoV-2 infection.

**Figure 4.**
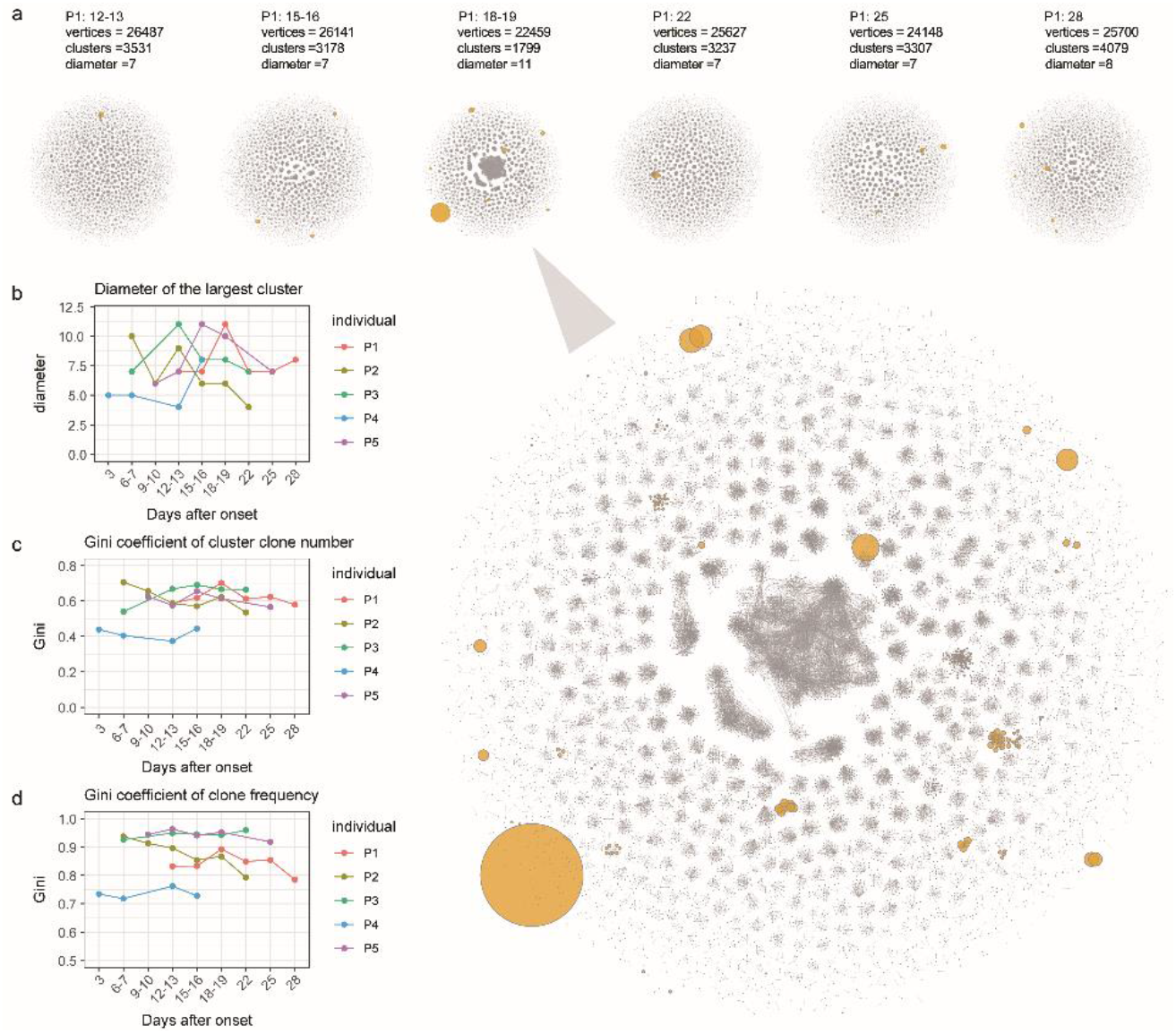
BCR repertoire network evolves over time after onset. **(a)** The network diagrams show how CDR3s are clustered with similar clones, each vertex represents a CDR3 amino acid sequence, the size of point represents the abundance of the clone, any two vertices meet three conditions, including same length, same V and J gene usage, and only one amino substitution are connected with edge. Six batches of network plot from individual P1 were lined up from left to right, the 18-19th day after onset were zoomed in. **(b-d)** cluster number, diameter of the biggest cluster and the average degree of the network, each dotted line represents the dynamic tendency of an individual over time after onset of the disease.

## DISCUSSION

To obtain a more detailed understanding of the dynamics and sophistication of B-cell immune response in COVID-19 patients, we applied a comprehensive analysis of IGH repertoire during the disease process. Given the advantages of longitudinal data in profiling the fast repertoire dynamics through time in individuals responding to an immune challenge ^[19, 24]^, we collected a set of longitudinal immune repertoire data from 5 COVID-19 patients and conducted a detailed analysis of their B-cell immune repertoires. The simultaneous profiling of clonotype overlap, lineage expansion and clonotype network enabled us to find that a period of 2-3 weeks after symptom onset is of great importance to B-cell immune response. In short, these results reveal a clear dynamic landscape of BCR repertoire in COVID-19 patients, as well as the characteristics of the immune repertoire at different time points after symptom onset.

V, J gene usage analysis revealed that each individual’s frequency of different V and J gene segment usage stays relatively stable over time (**Fig 1f**). Similar results have also been reported by Chen Wang et. al ^[6]^. Suggesting that V, J segment use is not markedly altered by SARS-CoV-2 infection. However, the overall use of V components among patients shows a differentiated preference. As Petter Brodin mentioned ^[22]^, this may reflect that the human immune system is highly variable between individuals but relatively stable over time within a given person. We observed a preference of the use of V gene segments inconsistent with the single cell BCR repertoire research by Wen et al ^[25]^, but consistent with Jacob’s ^[26]^ bulk BCR repertoire study, for example, Wen’s group reported that IGHV3-23 was over-represented in patients compared with health controls, while both our and Jacob’s result showed that IGHV3-23 is highly represented both in patients and healthy controls. Additionally, we and Jacob’s group both found that IGHV4-34 are more frequently used in COVID-19 patients compared with healthy donors. We believe that, given the advantages of large amounts of sequencing data, bulk BCR sequencing may be more suitable for BCR gene usage frequency assessment. Nevertheless, with only five patients, the power of our study is almost certainly constrained by a small sample size. Larger samples and more sequencing data need to be collected to help further understand these differences.

We found from the longitudinal immune repertoire data that all patients’ BCR repertoire composition in 2-3 weeks of onset is obviously different from that in other periods. There are significantly more common clones between adjacent time points and evident lineage expansion during 2-3 weeks. Additionally, network analysis revealed that each patient’s average degree and diameter indicators peaked around 2-3 weeks after onset. This timeline of repertoire change is consistent with an antigen (virus) induced B-cell differentiation and maturation. The maturation process is on a par with other viruses, such as influenza viruses ^[27]^, a similar phenomenon has also been observed in patients with Middle East respiratory syndrome (MERS) ^[28]^. We speculated that the higher clonotype overlap and substantial lineage expansion of COVID-19 patients during 2-3 weeks of illness may be associated with strong neutralizing antibody responses, accompanied by a large expansion of virus-specific B-cell clones and finally causing B-cell clone convergence.

Many studies have shown that the monoclonal antibodies targeting viral surface antigens have therapeutic and preventive effects on infectious diseases such as HIV, Ebola and MERS ^[29, 30, 31]^. Their safety and effectiveness in patients have been confirmed in many clinical trials ^[32, 33]^. With the efforts of scientists from various countries, recent positive progress has been made in the development of anti-SARS-CoV-2 drugs ^[34, 35, 36, 37, 38, 39]^. Our study can provide some clues in the development of neutralizing antibodies. The following rules should be taken into consideration in the screening of the SARS-CoV-2 neutralizing antibodies: Firstly, whether or not they are the highly overlapped clones during 2-3 weeks is of great potential. Secondly, the expansion size of clones which usually taken as an important criterion to select antibodies in other studies should also be involved. Subsequently, the cluster size in the network at the time window may also play their roles. It was interesting to find from the networks that highly expanded clones did not appear in the highly aggregated clusters, which was quite different from the blood cancer patients’ BCR repertoire network structure ^[23]^. We speculate that the big clusters are also important because they may be the B-cells iterating to gain a higher affinity to the virus antigen. Studies containing the B-cell isotype information may further reveal if the excessively clustered clones are more likely to be IgD/IgM naïve cells and the expanded clones are usually IgA/IgG effector cells. Nevertheless, the CDR3 sequence information from the selected antibodies may provide us the basis to produce specific drugs and vaccines.

In general, based on high-throughput immune repertoire sequencing technology, this study provides a comprehensive B cell deep immune profiling of COVID-19 patients, conclusions from this research will help understand the immune response of COVID-19 patients against SARS-CoV-2 virus and will provide the necessary basic knowledge for the development of vaccines and neutralizing antibodies.

## Acknowledgements

We sincerely thank the support provided by China National GeneBank. This study was supported by CAMS Research Units of Adaptive Evolution and Control of Emerging Viruses (2018RU009). William J. Liu is supported by the Excellent Young Scientist Program of the National Natural Science Foundation of China (NSFC, 81822040) and Beijing New-star Plan of Science and Technology (Z181100006218080). The study was supported by Guangdong Provincial Key Laboratory of Genome Read and Write (No. 2017B030301011), Shenzhen Science and Technology Innovation Fund (JCYJ20170412152916724 and JCYJ20170817145305022) and the COVID-19 Prevention and Treatment Fund from Shantou Science and Technology Agency. We thank Dr. Sunil Kumar Sahu at BGI-shenzhen for his linguistic assistance in this manuscript preparation.

## Data availability

The data that support the findings of this study have been deposited into CNGB Sequence Archive (CNSA: https://db.cngb.org/cnsa/) of CNGBdb with accession number CNP0001106.

## Author contributions

HX and LW analyzed the sequencing data and prepared the figures. YZ, PY, CrB, ZW, LY, JY, FM, CqB, NZ, YG, YB, SZ, and CZ collected the specimens, and retrieved and analyzed the clinical data. XL, MW, YZ, ZX, BY, PW and XL performed the experiments. HX, LW, MW and XL wrote the draft with the help of all authors. LT, NY, LL, YS, GW, WJL, YZ, PL and YG provided critical review of the manuscript. WJL, NY, PY and GFG conceived the idea and supervised the entire study. All other authors contributed to the work. All authors read and approved the manuscript for submission.

## Abbrevations

SARS-CoV-2: Severe acute respiratory syndrome coronavirus 2
COVID-19: coronavirus disease 2019
PBMC: peripheral blood mononuclear cells
BCR: B-cell receptor
IGH: immunoglobin heavy chain
CDR3: complementarity determining region 3
SHM: somatic hypermutation.

## Supplemental Figure and Table

**Supplementary 1.**
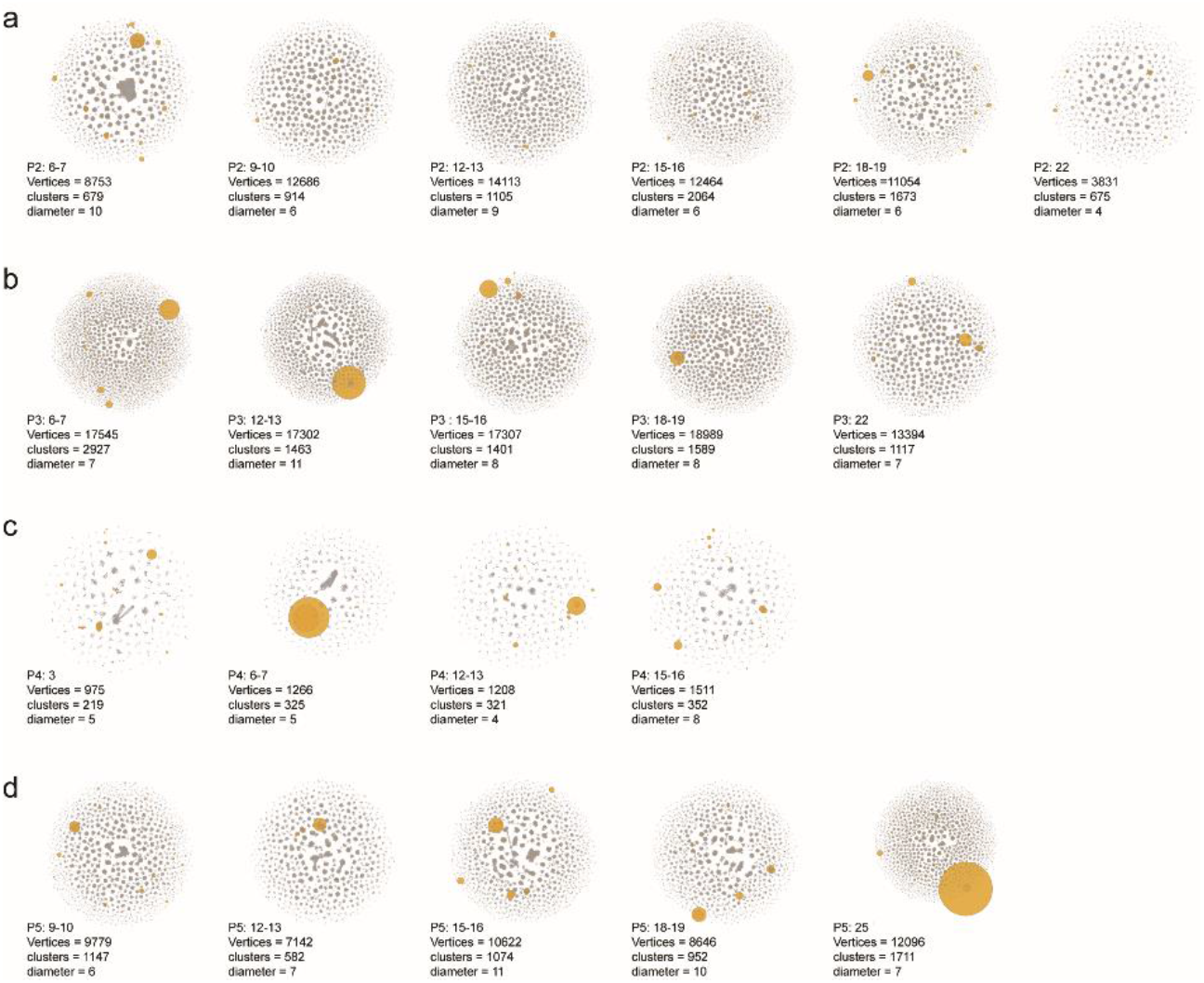
BCR repertoire network evolves over time after onset. The network diagrams show how CDR3s are clustered with similarity, each vertex represents a CDR3 amino acid sequence, the size of point is proportional to the abundance of the clone, any two vertices meet three conditions, including same length, same V and J gene usage, and one amino substitution are connected with edge. (a-d) Different batches of BCR repertoire network from individual P2 to P5 are lined up from left to right.

**Supplemental Table 1.**
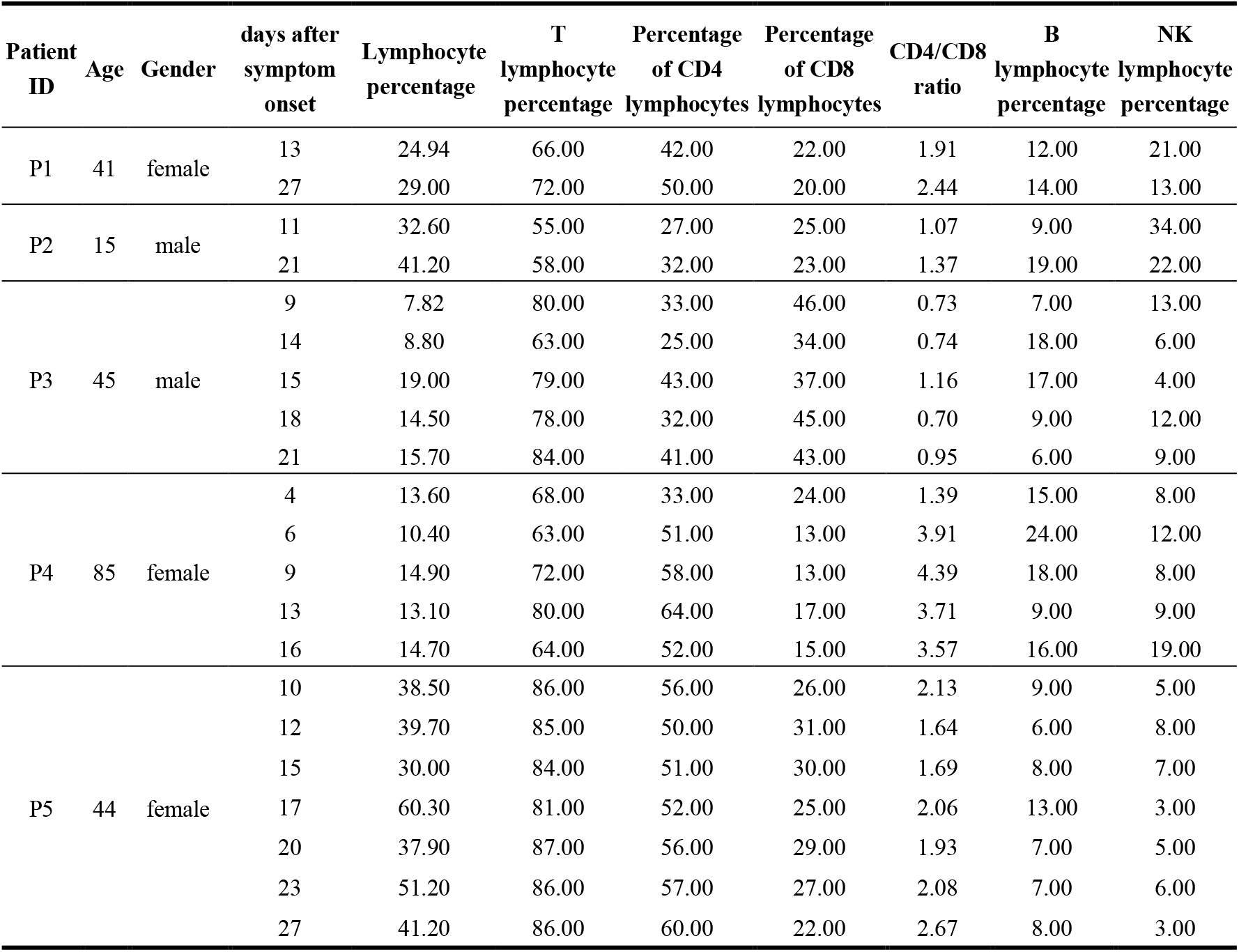
Summary of the laboratory characteristics of the 5 convalescent patients.

